# A method for assessing approach and avoidance behavior across multiple olfactory stimuli in mice including multivariate hypothesis comparisons

**DOI:** 10.64898/2026.02.21.707096

**Authors:** Michelle C. Rosenthal, Alper Kagan Bakir, Ankur Gaikwad, Katharina Clark, Adam T. Garcia, John P. McGann

**Affiliations:** Behavioral and Systems Neuroscience Department of Psychology; Rutgers Center for Cognitive Science Rutgers University New Brunswick; Department of Otolaryngology-Head and Neck Surgery, Rutgers Robert Wood Johnson Medical School; Rutgers, The State University of New Jersey

**Keywords:** generalization, fear, perception, smell, hypothesis-testing, open field

## Abstract

Approach and avoidance behavior towards sensory stimuli serve as powerful behavioral readouts of our mental representations of the external world and our expectations and motivations in navigating it. In the olfactory system, approach or avoidance of odors statistically associated with people, places, and things relate to ecologically critical functions like feeding, fear, and reproduction. However, experimental methods for quantifying approach/avoidance behavior in relative terms across odors have been limited. Here we present a novel method for quantifying mouse approach/avoidance in an open field arena scented with up to four odors simultaneously. In lieu of traditional inferential statistics (which greatly limit the information that can be learned in this multivariate experiment), we demonstrate the *a priori* definition of quantitative hypotheses for the distribution of time among scented corners and the use of information theory-derived statistical metrics to quantify the relative likelihood of each competing hypothesis given the data collected. Finally, we use data from a fear conditioning experiment to demonstrate the application of this method to conclude that fear conditioned mice exhibit a fear generalization gradient that decreases as odorants become more different from the threat-predictive odorant, as opposed to competing hypotheses that mice are specifically avoiding the threat-predictive odorant or have overgeneralized their fear and avoid all test odors regardless of similarity. Critically, this method takes only a few minutes per animal with no prior behavioral training required, and it can be performed easily without automated apparatus.

## Introduction

Stimulus avoidance and its complement stimulus approach played a critical role in early theories of mind and behavior, including models of personality, mental health disorders, and traumatic stress (Lewin 1935; Miller 1944; Solomon *et al*. 1953). Avoidance behaviors have thus been among the most important dependent measures in psychological science, including for the investigation of perception, learning, and behavioral disorders. In animal models, avoidance is typically assessed via place preference testing, in which the location of a freely moving animal relative to one or more test stimuli is quantified over a modest period of time in a shuttlebox (Mathur *et al*. 2011; Mowrer and Miller 1942; Scherma *et al*. 2021), open field arena (Hall and Ballachey 1932; Kulesskaya and Voikar 2014), or similar apparatus. Usually this is an assessment of passive approach/avoidance behavior, though sometimes more complex contingencies are incorporated (Harlow 1958).

It is common in learning and memory experiments to test how specific or general a memory is by training a subject to make a response when presented with a particular stimulus (often called the conditioned stimulus, or CS) and then testing whether the subject responds to each of a succession of different stimuli related in some way to the CS (Lashley and Wade 1946; Pavlov 1927). When a subject trained on a CS also responds to non-conditioned stimuli, this is referred to as *generalization* from the CS to the test stimulus (Lashley and Wade 1946). Most commonly the stimuli are varied along sensory dimensions, such as stimulus magnitude, color, or pitch (Dunsmoor and LaBar 2013; Guttman and Kalish 1956; Laxmi *et al*. 2003). In most cases the test stimuli that most resemble the CS are the ones that exhibit the most generalization, with decreasing similarity to the CS yielding decreased responding. This fall-off of generalization with decreasing similarity is called a *generalization gradient* (Dunsmoor *et al*. 2017; Dunsmoor and LaBar 2013; Guttman and Kalish 1956). Differences in generalization gradients between subjects or experimental conditions have been the basis of landmark studies in the exploration of human and animal behavior (Guttman and Kalish 1956; Hanson 1957; 1959; Harlow 1958; Murray and Miller 1952). Classical generalization designs potentially confound differences in sensory discrimination (what does the subject think the new stimulus is) with differences in stimulus-response coupling (what outcome does the subject think the new stimulus predicts). However, generalization of learning can be decoupled from perceptual ambiguity by sensory training (Hanson 1957; 1959) and generalization occurs along non-sensory dimensions like conceptual similarity (Dunsmoor *et al*. 2011; Fleurkens *et al*. 2011; Gerdes *et al*. 2020), thus establishing that generalization of learning is not merely a sensory phenomenon.

Generalization of fear learning to new stimuli is adaptive, but overgeneralization of fear to stimuli and situations that do not in fact predict a threat is a potential mechanism of anxiety disorders in humans, including generalized anxiety disorder, panic disorder, and posttraumatic stress disorder (Dunsmoor *et al*. 2017; Dunsmoor and Paz 2015; Fraunfelter *et al*. 2022; Lissek *et al*. 2010). Fear generalization following conditioning with olfactory stimuli produces diverse effects on olfactory processing, including changes in early olfactory signal processing (Chen *et al*. 2011; Kass and McGann 2017; Pavesi *et al*. 2012; Ross and Fletcher 2018b), that are different than the changes evoked by learned fear of a specific odor (Ahs *et al*. 2013; Kass *et al*. 2013b; Li *et al*. 2008; Li *et al*. 2006; Parma *et al*. 2015).

Conversely, human subjects with high levels of trait anxiety exhibit increased generalization between odors in a discriminative aversive conditioning paradigm and fail to improve their perceptual discrimination between the odors (Rosenthal *et al*. 2022). There is thus a specific need for improved methods for evaluating fear generalization in the olfactory system.

Stimulus generalization testing is typically performed in trial-based learning tasks and thus uses sequential presentations of individual stimuli (or occasionally pairs of stimuli such as in a two-lever choice paradigm) rather than simultaneous presentation of multiple stimuli (Ross and Fletcher 2018a; Youngentob *et al*. 1990). This has served both the theoretical purpose of ensuring that the behavioral response to each stimulus is assessed independently without confounding effects of stimulus contrast and the practical purpose of permitting hand scoring the position of the subject with a stopwatch in the analog era. However, sequential presentations incur new learning as each stimulus goes unreinforced, which confounds the original stimulus-response relationship being tested and puts a limit on the number of new stimuli that can be presented to a single subject. Here we demonstrate a combination of spatial place preference testing with hypothesis testing that enables simultaneous testing of four simultaneous stimuli, thus limiting the effects of presentation order and new learning.

Modern computerized video analysis has pushed the spatiotemporal resolution of place preference data to its practical limit, with the latest methods able to extract profound insights about the microstructure of the animal’s behavior (Lauer *et al*. 2022; Mathis *et al*. 2018; Weinreb *et al*. 2024; Wiltschko *et al*. 2015). However, this resolution has paradoxically created new challenges for interpreting the complex and idiosyncratic behavior of individual subjects relative to *a priori* hypotheses. Introducing multiple stimuli to a place preference task greatly complicates classical approaches to statistical hypothesis testing because rejecting the null hypothesis (a uniform distribution of subject position in space) sheds little light on the actual approach/avoidance behavior of the subject. Moreover, frequentist approaches based on p-values lose power rapidly as the number of variables increases because of the need to correct the alpha value for number of comparisons (Benjamini 2010; Miller 1981). Here we demonstrate how to develop *a priori* quantitative hypotheses of how different generalization processes would manifest in different distributions of mouse positions. Then we show how to apply the likelihood ratio, which is the uniform most powerful test for this context (Neyman and Pearson 1933), and the Akaike Information Criterion (AIC) (Akaike 1974) to compute the relative likelihood of each hypothesis given the behavioral data collected.

### Behavioral testing

The open-field setup consists of a custom-built 18×18-inch box with 18-inch high white matte walls and floor (Fig. 1A) that was novel to the mouse during testing. A 2-inch round metal mesh tea strainer was suspended in each corner to act as a local odor diffuser (Fig. 1B). As in training, each tea strainer contained a 0.5 inch diameter cotton ball treated with 200 microliters of pure odor (or no odor as a control) immediately before testing each animal. Each mouse is placed in the center of the box and allowed to explore freely for 5 minutes, after which they were returned to their individual cages. Mice were continuously recorded using a fish-eye lens (Computar 4.5-12.5mm) and camera (Basler 106558). In some experiments not reported here we have used an alternative commercially available arena that automatically quantifies mouse position through infrared beam breaks (Kinder Scientific, Poway, CA), but this apparatus did not meaningfully improve the resulting behavior quantification.

**Figure 1.**
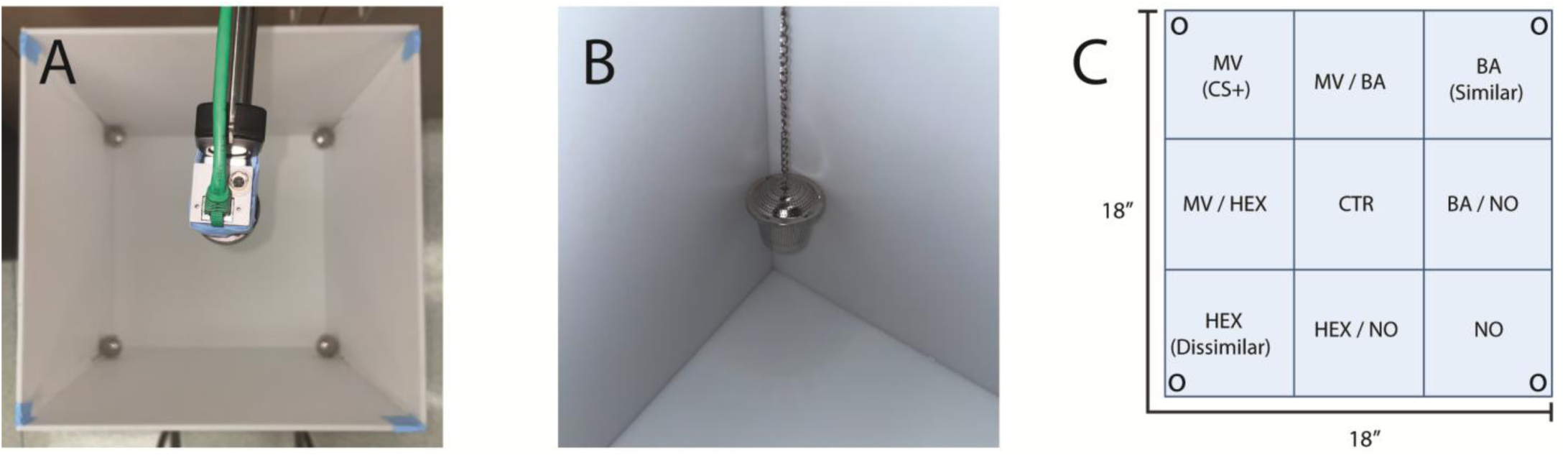
Behavioral apparatus. Mice are placed in a square, high-sided arena with a camera overhead (A) and tea diffuser balls containing odorant suspended in each corner (B). For the fear generalization testing reported here, the odors were laid out as shown (C), with the odor that predicted shock (MV, the CS+) and the No Odor corner (NO) opposite each other. For analysis, the position of the mouse was assigned to one of the nine zones depicted (C). Blue tape was used in A to illustrate how tea balls were suspended, but identical pieces of white or transparent tape are recommended during use. *Alt text: Photographs of the arena and odor diffusers in subpanels A and B with a schematic of arena scoring zones in subpanel C*.

The odors included methyl valerate (MV, a somewhat fruity ester odor), butyl acetate (BA, another ester that smells less fruity), 2-hexanone (Hex, a ketone that smells quite different from the two esters), and no odor. The assignment of the odors to corners is a critical part of the design of this experiment: the shock-predictive odorant goes in one corner and the no odorant control goes in the opposite corner. The other two odorants thus provide equidistant options from both safety and threat. We typically use methyl valerate as the threat-predictive stimulus and see a generalization gradient where MV-conditioned mice spend more time in the No Odor and Hex corners (and the “safe” zone between them) and less time in the MV and BA corners and the “danger zone” between them). To counterbalance the locations of odors within the room, the arena was rotated from animal to animal. The arena is positioned so that each corner is equally lit by room lighting. The tea strainers are cleaned with ethanol and rinsed with water in between animals.

We usually use esters (e.g. MV, BA, ethyl tiglate, isoamyl acetate, etc.) and ketones (e.g. 2-hexanone, 2-heptanone) as odorants for these experiments because they preferentially drive neural activity in the region of the dorsal olfactory bulb visualized during our optical neurophysiology experiments (Fast and McGann 2017; Kass *et al*. 2013a). Alternative sets of odorants can be chosen to match the needs of other experiments.

In selecting sets of odorants for experimental use it is common to consider issues of odor chemistry and perception. All odorants differ in their vapor pressure and thus achieve different concentrations in the air in an open container. It is thus useful to select odorants with similar vapor pressures. We have sometimes corrected for differences in vapor pressure between odorants by proportionately diluting the higher vapor pressure odorants in mineral oil (Jennings *et al*. 2023). In experiments that use odorant concentration as a cue this adjustment could potentially be important, but in the experiments presented here the concentration is not predictive of threat. In experiments with human subjects it is common to adjust odorant deliveries not to match the chemical concentrations achieved but to equate perceptual features like intensity or pleasantness (Cain 1971; Svensson and Lindvall 1974). In mice these metrics are not directly accessible. Despite these potential complexities, we find that in practice unconditioned control mice really do distribute their time around the arena without regard to the odorants at the concentrations employed here (see below), suggesting that any differences in concentration/intensity or pleasantness are not sufficient to influence the assessed behavior. However, we caution that higher concentrations of these odorants could be inherently aversive to mice, as could certain other odorants that mice find inherently aversive (Francesconi *et al*. 2020). We also note that some odorants can interact with each other in the nose through competitive antagonism and allosteric modulation (de March *et al*. 2020; Inagaki *et al*. 2020; Xu *et al*. 2020), which could impact perceptual quality in regions of odor overlap.

### Behavior quantification

Data in the arena may be quantified using any number of methods, including manual scoring, use of commercial software (e.g. EthoVision), or use of open source pose estimation tools (Mathis *et al*. 2018). We used EthoVision to compute two main measures: time spent in each zone and the average distance from specific corners. Due to slight changes in the arena orientation due to counterbalancing and each animal being of different size and shape, each trial had its own pair of arena and detection settings. After selecting the arena’s external boundaries within the field of view, the open-field arena was then divided into 9 equal square zones, including odor zones, intermediate zones, and the center.

Unequal-size zones could conceivably be useful in other experiments, in which case the measured dwell times would be corrected by the area of each zone to make a time density metric. Each odor corner was marked as a point in the EthoVision software for distance calculations. The time spent in each zone and mean distance were determined based on the animal’s center point in the video. The animals were identified using EthoVision’s automated method for each animal specifically, this detection setting was then manually tested to make sure it is properly identifying the animal. Ethovision’s mobility score, which represents the fraction of the time that the mouse was classified as moving, was defined as the total interframe intervals in which a frame-to-frame comparison showed the body area occluding a different part of the arena floor (the threshold is determined dynamically by EthoVision based on the optical contrast between each mouse and the arena floor). We note that while EthoVision was a convenient and available tool, especially for providing moment-to-moment measurements like mobility and real-time distance between the mouse and each stimulus, in principle the key metric of time spent in each zone could be measured manually during video playback using a simple timer. All statistical analyses were performed in RStudio, and graphs were created using RStudio, Python and Adobe Illustrator.

### Quantitative hypothesis specification

Modeling of data has two general approaches: fitting of *a priori* models as hypothetical explanations of the data and fitting of *a posteriori* models to characterize the underlying structure of the data as it is. Here we focus on *a priori* quantitative models, which can be thought of as precise, competing hypotheses about how subjects might behave. We emphasize that the goal is to produce different, conceptually contrasting models, not necessarily to figure out the best possible fit to the data actually collected. We recommend stating the model in words and then working through the logic to define the actual numbers. For example, for the generalizing fear conditioning experiment below we verbally described the following *a priori* models (Fig. 2):

- **No Fear Model**, conceptualized as equal time spent in all areas (a.k.a. chance model)
- **Fear Without Generalization Model**, conceptualized as minimal time spent in the CS zone, 50% reduced time in the CS-adjacent sides, and the balance of time distributed uniformly across all other regions
- **Fear Overgeneralization Model**, conceptualized as minimal time spent in all scented corners and the sides between them, 50% reduced time in the sides between an odor and the No Odor corner, and the balance of time distributed uniformly across all other regions
- **Gradient Fear Model**, conceptualized as minimal time spent in the CS-zone, reduced time in the other scented corners in proportion to the similarity to the CS, reduced time in the sides between the CS odor and the other odors that is average of the dwell time in each corner, and the balance of time distributed uniformly across other regions.

**Figure 2.**
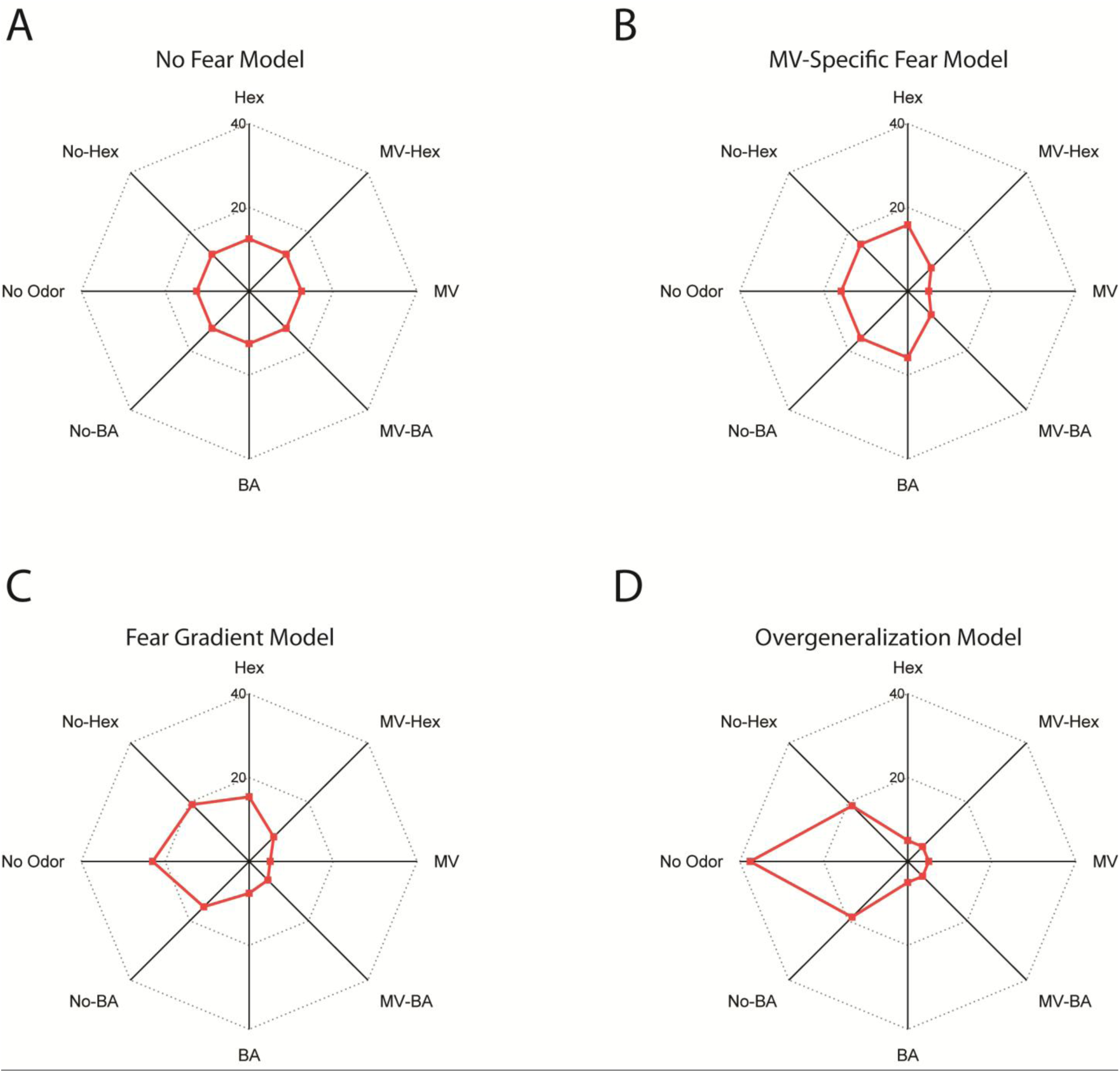
*A priori* models of potential fear generalization behavior. These radar plots depict the proportion of time in the arena spent in each zone (from 0 to 40%), where the corners are scented with individual odors (MV, BA, HEX, or No Odor) and each side of the arena is denoted by the pair of corners it connects (e.g. MV-HEX). Models depict the four *a priori* hypotheses: No Fear (A) in which the mouse spends equal time in each zone; MV Specific Fear (B), in which the mouse is only afraid of MV and spends the minimum time (5%) in the MV zone and reduced time in the MV-adjacent zones; Fear Gradient (C), in which the mouse is afraid of each odor in proportion to the similarity of its neural representation to that of MV and thus avoids MV most, then BA, then Hex; and Overgeneralization (D) in which the mouse is equally afraid of all three odors and spends the most time in the No Odor corner. *Alt text: Figure showing four graphs visually depicting the a priori models of time distribution around the arena*.

Instantiating these models in practice requires selecting some additional constraints:

- *Minimum dwell times:* We define a non-zero minimum for each zone, since mice generally enter each zone at least briefly, possibly to identify the stimulus. Even mice that are highly afraid of the CS usually spend at least a few seconds in the CS corner, so we use a minimum of 5% time (15 sec) in any zone in constructing our hypothetical models. This minimum was selected based on previous real-world data on this task and might reasonably be tuned *post-hoc* to update the hypotheses to match the actual minimum dwell times obtained.
- *Treatment of center zone:* We have elected not to include the center zone in our model, but to focus on the relative distribution of time around the periphery of the arena. This reflects the nature of our hypotheses, which are about the avoidance of odors relative to each other, rather than hypothesizing about the position of the mouse relative to the center of the maze. It also makes the model distribution circular (matching the perimeter of the arena), where traveling in one direction eventually brings you back to where you started. If formulating a hypothesis that relates to the preference to be against a wall instead of in the open middle of an arena or treats all corners identically, then including the center zone would not be a problem. Empirically, our fear conditioning experiments have all produced about the same percentage of time spent in the center across groups. If excluding the center zone, the data from the other zones should be normalized (scaled so that the total is 100%) prior to fitting the models. Note that in principle one could alter the physical arena to make it annular rather than square, thus eliminating the center zone, but that a) eliminates the neutral starting point, and b) forces subjects to traverse other zones to get to their preferred zone.
- *Treatment of arena sides*: In principle it is a decision to structure the model on the presumption that dwell time in the side zones of the maze will reflect the average of the dwell times in the adjacent corners. Alternative models are possible, such as hypothesizing that the sides are more like the No Odor corner than like their adjacent odors (since they are farther from the odor source) or hypothesizing that a side between an odor corner and an unscented corner will act more like one or the other. However, averaging is the most parsimonious model and empirically works for the odor concentrations employed here. In principle one could exclude the side zones on the same logic as excluding the center zone, but we find that mice sometimes spend quite a lot of time in these zones and that such time can be very informative.

Converting conceptual hypotheses to the quantitative hypotheses in Table 1 can be more challenging than it appears. We recommend the following steps:

1. Plug in the known values. For instance, the MV Fear Without Generalization model specifies the minimum 5% dwell time in the MV corner, while the Overgeneralization model specifies the minimum 5% dwell time in all odor corners and sides between odor corners.
2. Compute the total share of time left to allocate after the known values are accounted for. For the MV Fear without Generalization model this is 95%, while for the Overgeneralization model it is 75%.
3. Assign relative values to all remaining corners. For instance, in the MV Fear Without Generalization models, the BA, Hex, and No Odors would all be the same, so we assigned them each 1 share of the unallocated time. The Gradient fear model by contrast assigned values of 0.33, 0.67, and 1 to those corners based on neurophysiological similarity data.
4. For any still-unspecified side zone, calculate the proportional share as the average of the shares of the adjacent corners.
5. Divide the unallocated time from step 2 by the total number of relative shares from steps 3 & 4 to get a metric of time spent per relative share.
6. Multiply the relative shares of each zone from steps 2 & 3 by the time per share calculated in step 5 to get the final values for each zone of the model.
7. Total the percentages across the 8 zones to confirm they account for 100% of the time.
8. Graph the results to visualize the hypotheses (Fig. 2). We prefer radar plots (available in virtually all graphing software including Microsoft Excel) because they intuitively capture the multidimensional aspect of the data by recapitulating the physical relationships among the odor zones in the arena and because they can show error bars if desired to show confidence. A chance distribution of time across eight zones will make a perfect octagon (Fig. 2A), while deviations from

**Table 1.** Four *a priori* models described by their hypothesized relative dwell time in each zone as a percentage of the total time in the perimeter zones. Gradient Fear model is calculated presuming a 67% similarity between MV and BA and a 33% similarity between MV and HEX, numbers chosen to approximate their relative similarity based on chemical structure (ester vs. ester compared to ester vs. ketone) and published glomerular response maps.

|  | No Fear | MV-Specific Fear | Overgeneralization | Gradient Fear |
| --- | --- | --- | --- | --- |
| Hex | 12.5% | 15.83% | 5% | 15.4% |
| MV-HEX | 12.5% | 7.917% | 5% | 8.3% |
| MV | 12.5% | 5.00% | 5% | 5% |
| MV-BA | 12.5% | 7.917% | 5% | 6.3% |
| BA | 12.5% | 15.83% | 5% | 7.6% |
| No-BA | 12.5% | 15.83% | 18.75% | 15.3% |
| No Odor | 12.5% | 15.83% | 37.5% | 23.0% |
| No-Hex | 12.5% | 15.83% | 18.75% | 19.2% |
| Total | 100 | 100 | 100 | 100 |

### Visualizing data and quantifying the fit of each hypothesis

Generally, the first step in fitting hypotheses to data is to visualize the quantified data. Many commercially available systems can convert position data into heat maps, which is a useful first pass visualization of data from an individual subject. While we are well aware of the dangers of averaging individual subjects, in practice we have found that if the data is quantified into nine zones as described above, averaging those nine measurements across subjects produces a reasonable summary of group behavior. Often comparing the group data plots to the hypothesis plots immediately reveals one model that is obviously the best fitting, but we nonetheless advise a quantitative assessment of fit.

In principle, there need be no actual “fitting” of each hypothesis to the data because both are precisely defined, so a simple error term can be precisely calculated. However, in practice we recommend the inclusion of a single scaling parameter, applied equally throughout the zones, to ensure that the model is testing relative time across the scented zones. It also allows the user to harness existing software tools that are designed for fitting custom functions. If the data is not normalized (e.g. if different total times are possible across animals, such as from excluding time in the center zone) then this scaling parameter is required. The key step in fitting the hypotheses is the expression of each hypothesis as a mathematical function. For most hypotheses, including all except the No Fear model detailed above, this function will be discontinuous such that each zone has its own value different from the adjacent zones on either side. The hypotheses are thus represented as *piecewise functions*, in which the rule for each subdomain (i.e. each zone) is different from the others. The form of the function for the hypotheses described above is f(x)=ay_i_, where x is the number of the zone (1 through 8 in any order), y is the hypothesized value, and a is the scaling parameter.

Many statistical programs enable the user to define and fit piecewise functions to data, or it can be done in coding packages in R. Fig. 3 shows a block of C code that we used in Origin Pro software (OriginLab, Northhampton, MA) as a User Defined Function called nlsfMVonlyFear_floor to illustrate the use of a “switch” statement to easily define the discrete mapping of zones within the function. A series of if/else statements could serve instead and might better accommodate some hypotheses. This function is then fitted to the behavioral data using the existing tools in most statistical software, typically a curve fitting package based on the Levenberg-Marquardt gradient descent algorithm. Note that the data points can be “weighted” in curve fits so that some have a stronger influence than others. When fitting data that is an average of multiple subjects, we recommend that the inverse of the standard error of the mean (i.e. 1 divided by standard error) be used as the weight parameter for each zone. We also note that it is generally necessary to provide the value 1 as an initial value for parameter a. We have never had a properly configured hypothesis fit fail to converge using this approach, but we have frequently experienced convergence failure upon forgetting to properly initialize parameter A.

**Figure 3.**
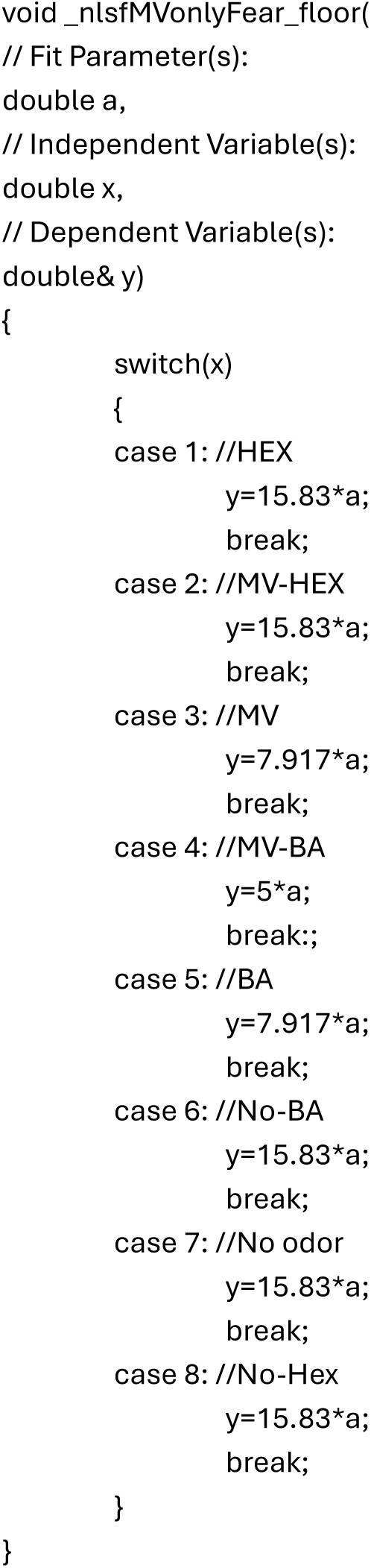
Example code block defining the discontinuous function instantiating the “MV Specific Fear hypothesis” in C. The values of x represent the different zones around the perimeter of the arena, beginning with the No Odor-Hex side zone and proceeding to the Hex zone. *Alt text*: *A block of computer code implementing a discontinuous function*.

### Computing AIC and relative likelihood among hypotheses

Having fitted multiple competing hypotheses to the data, it is then necessary to choose among them. There are multiple ways to make this evaluation, but we recommend using the Akaike Information Criterion (AIC). Briefly, the AIC computes the Kullback-Leibler divergence between the behavioral data and the fitted model to produce a metric of the information that would be lost by replacing the actual data with the model of the data (Akaike 1974). Lower values of the AIC thus indicate a better fit. The AIC metric enables model selection among disparate models (without any requirement that the models be nested) by computing the relative likelihood of one model over another using the formula:

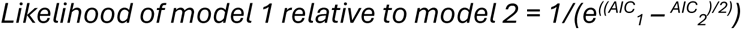

On the basis of this ratio, it is possible to quantitatively state how much more likely the closest hypothesis is than the second-closest hypothesis (Neyman and Pearson 1933). If a chance model (equal time in all zones) is included among the hypotheses, then it is also possible to quantitatively state how much more likely the best fitting model is than random chance. This likelihood metric is an alternative to both frequentist and Bayesian approaches to statistical inference (Neyman and Pearson 1933). However, it is important to note that because only *relative* likelihoods are computed it is possible to have a model that is much more likely than others and yet nonetheless not actually a good descriptor of the data.

Direct visual examination of the fitted function compared to the underlying dataset is recommended, as is consideration of primary goodness of fit metrics like chi-square or r^2^ (the coefficient of determination) and review of the residuals.

### Demonstration: Behavioral assessment of odor-cued fear generalization

#### Design

To demonstrate the utility of this methodology for assessing the generalization of fear across odors, we trained mice using a context-like fear conditioning paradigm (Bakir *et al*. 2026), then tested their fear generalization based on the methods described above. Briefly, the fear conditioning paradigm consisted of three 10 minute, once-per-day pre-exposures to the conditioning chamber (Coulbourn Apparatus) without odors or shocks, then three 10 minute, once-per-day conditioning sessions in which the chamber was scented with 200 methyl valerate (MV) placed on a cotton ball below the chamber. Mice in the experimental group (fear conditioned group, N=20) received 10 mild shocks (0.3 mA, 1 sec duration) randomly timed to occur every 40-80 seconds. Mice in the control groups received the same conditioning paradigm but with either the shocks omitted (Odor alone group, N=12) or the odors omitted (Shock alone group, N=12). Mice were then tested for fear generalization in the odor arena. As shown below, mice in the Odor alone group and Shock alone groups behaved identically during testing and were pooled into a combined Control group (N=24) for subsequent analysis and data display.

#### Mice

The demonstration data used a total of 44 C57BL/6 strain mice aged between 3 and 9 months and obtained from Jackson Laboratories. Mice were housed under a 12-hour light/dark cycle with *ad libitum* access to food and water. These demonstration data were drawn from control groups from behavioral pharmacology experiments (Bakir *et al*. 2026), and so all mice were fitted with bilateral olfactory bulb cannulae prior to the experiment and infused with a saline vehicle prior to testing. The animals were otherwise experimentally naïve. We have observed similar results in surgically naïve mice in other studies. To prevent injury, mice were housed individually following cannula implantation. The work was performed in compliance with protocols approved by the Rutgers University Institutional Animal Care and Use Committee (IACUC).

## Results

Comparing the data from the control mice and fear conditioned mice revealed modest changes in overall locomotion, with fear conditioned mice exhibiting a moderate but significant reduction in time spent moving (Fig. 4A) compared to control animals (t_42_ = 2.02, p = 0.0498, Cohen’s d = 0.6). Fear conditioned mice also spent moderately but significantly more time in the 8 boundary zones than the center zone (Fig. 4B) compared to the control animals (t_42_ = 2.74, p = 0.009, Cohen’s d = 0.81). This combination of reduced mobility and heightened thigmotaxis (preferring a wall over an exposed center region) is consistent with odor-cued fear in the conditioned mice but does not provide any information about the odor-specificity or generalization of that fear.

**Figure 4.**
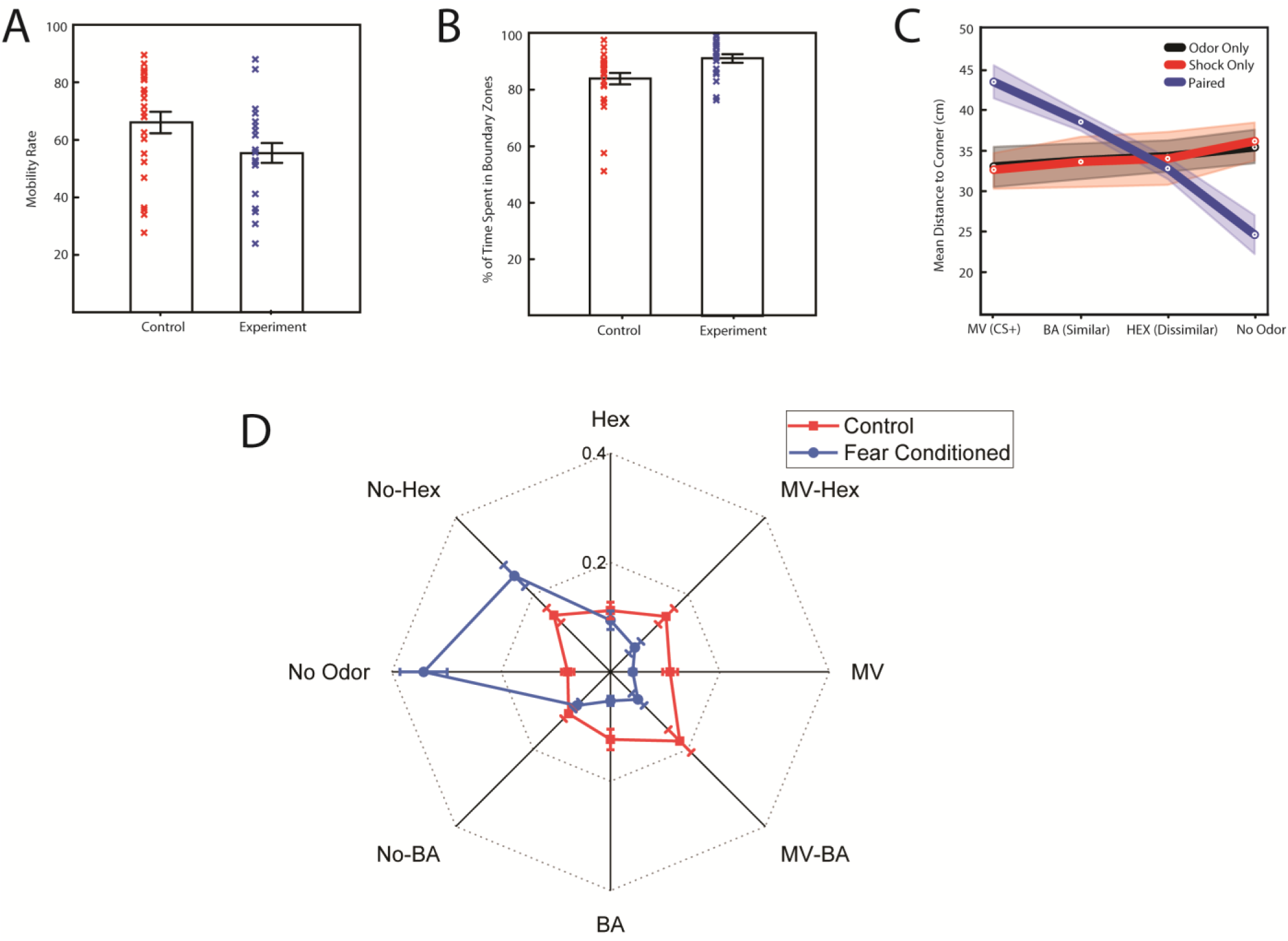
Fear conditioning results. Mice that underwent fear conditioning (blue) spent slightly less total time exploring the arena (A) and slightly more time in the boundary zones than in the center of the arena (B) compared to mice from the non-conditioned control groups (red). The position of fear conditioned mice (C, blue line) in the arena was on average closest to the No Odor corner, with successively more distance from the Hex corner, BA corner, and MV corner. Control groups receiving odor alone exposures or shock alone exposures were comparably far from all four corners on average (C, red and black lines) and nearly identical to each other. Color bands around each line illustrate the standard errors of the mean. The time spent in each boundary zone is shown in D, with fear conditioned mice (blue) distributing their time towards the No Odor corner and No Odor-Hex side, with much less time spent in the MV and BA corners and the corresponding sides. Control mice (red) distributed their time fairly equally among the zones. *Alt text: Graphs depicting mobility, distance to the corners, and time spent in each zone of the arena for control and fear conditioned mice*.

As an objective measurement of avoidance, we used EthoVision to calculate the distance of the center of the mouse to the odor stimulus in each corner of the arena on each frame of the video. Fig. 4C shows the average distance from each of the corners across the five minute session for each experimental group. The odor alone (black) and shock alone (red) control groups both average an equal distance from each corner, producing a horizontal line on the graph. Note that the two lines perfectly overlap, showing that these two groups did not differ from each other. By contrast, the fear conditioned mice (which received Paired MV-shock training) were on average closest to the No Odor corner and furthest from the MV corner (Fig. 4C, blue). Note that this distance metric also suggests a generalization gradient, in that the fear conditioned mice (blue) stayed farther away from BA (a similar, yet readily distinguishable odor from MV) than controls did (red and black), yet did not stay away from Hex (a more different odor). These data can be analyzed statistically in many ways, including both linear and non-linear models, Bayesian frameworks, and other approaches, but traditional frequentist approaches are complicated by the presence of four dependent variables (distance to each corner) that are not independent of each other. For simplicity, we note that the traditional frequentist approach to avoidance reveals a significant effect of training group on distance from MV (one-way ANOVA, F_2,41_ = 29.05, p < 0.001, eta-squared = 0.59), with the Paired group significantly larger than the Shock Only group (p < 0.01, Cohen’s d = 2.38) and the Odor Only group (p < 0.001, d = −2.17). However, it would be difficult to correctly use a method like ANOVA to statistically demonstrate that these data reflect a generalization gradient, even with these robust results, because of the need to correct the alpha value for multiple comparisons. These distance numbers also only implicitly capture the positioning of the mouse, leaving it ambiguous whether the mouse is in the center, the corners, the sides, etc.

A yet richer picture emerges when we analyze the time spent in each boundary zone and display them on an eight-dimensional radar plot corresponding to the odor layout of the arena (Fig. 4D). This display, which is analogous to a heat map, provides an intuitive display of where mice actually went in the arena while also displaying cross-animal variance via error bars. As shown in Fig. 4D, non-conditioned mice in the Control group spent similar amounts of time in all eight boundary zones of the arena, while the fear conditioned animals reduced their time in the MV, MV-BA, BA and MV-Hex zones and extended their time in the No Odor corner and neighboring No Odor-Hex side. It allows us both to make easy comparisons (e.g. the fear conditioned mice spend less time than controls in the BA corner but not the Hex corner) and to recognize overall patterns (the mice went towards the Hex side of No Odor), but it does not by itself enable us to make quantitative claims about those patterns.

The final step is to compare the observed behavioral data to the *a priori* models described above. Fig. 5A displays the No Fear model (red) superimposed on the behavioral data from the control group mice (blue), which clearly matches the data well. Computing the actual fits between the control data and the four models described above confirms this visual impression (Table 2), with the No Fear model matching the empirical distribution with modest residuals and a low AIC. Unsurprisingly, the No Fear model is thus 100-fold more likely than the MV-Specific Fear model and orders of magnitude more likely than the other two models (Table 2). In contrast, Fig. 5B displays the Fear Gradient model (red) superimposed on the empirical data from the fear conditioned mice (blue). This model matches the data reasonably well but not perfectly, with some evidence that the mice might be somewhat more afraid of Hex and BA than the model envisioned. This is precisely the situation where comparative modeling is valuable. As shown in Table 3, the Gradient Fear model produces the best fit, with the smallest residuals and AIC. Based on the relative likelihoods of Gradient Fear model and the competing models (Table 3), we can draw a series of conclusions: a) the mice are afraid of at least one odor (24-fold likelihood vs No Fear), b) mice are afraid of more than just MV (11-fold likelihood vs MV-Specific Fear), and c) mice are not equally afraid of all odors (3.4-fold likelihood vs Overgeneralization), though that is the weakest of the three claims. This demonstrates the utility of this hypothesis testing approach over the more traditional analysis methods above.

**Figure 5.**
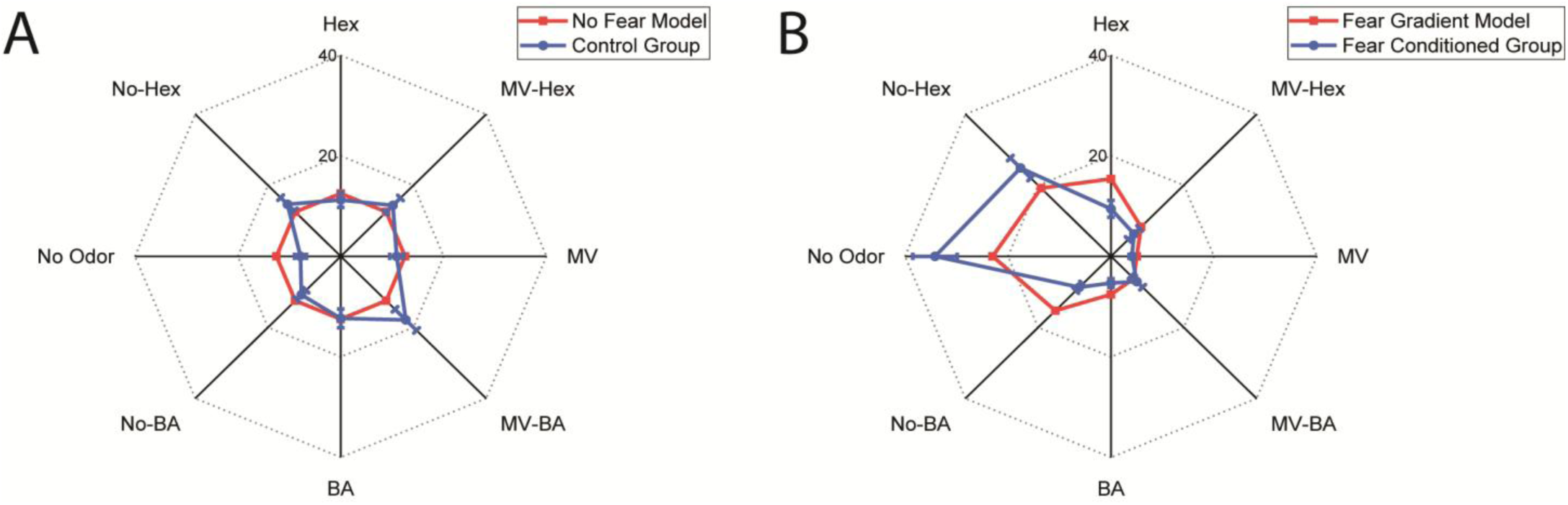
Fits to empirical data. A) Plot of time spent in each boundary zone in the control (unconditioned) groups (blue) compared to the *a priori* No Fear model. B) Plot of time spent in each boundary zone for the fear conditioned group (blue) compared to the Fear Gradient model (red). Error bars represent one standard error. *Alt text*: *Graphs depicting the empirical distribution of time in each zone compared to the hypothesized distribution of time in each zone*.

**Table 2.** Model fits for Control Mice. Top table shows the quality of the fit of each model to the behavioral data, expressed as residual sum of squares (RSS, lower shows a better match between model and data), and Akaike Information Criterion (AIC, lower indicates a better match between model and data). Lower table shows the likelihood ratio between the best fitting model (No Fear) and the other models based on their AIC scores.

| Model (Control Group) | RSS | AIC |
| --- | --- | --- |
| No Fear | 19.9 | 13.70 |
| CS-specific fear | 63.0 | 22.91 |
| Gradient Fear | 131.8 | 28.82 |
| Overgeneralization | 232.6 | 33.36 |

**Table 2. Model fits for Control Mice.**
| Comparison (Control Group) | Likelihood Ratio (fold likelihood) |
| --- | --- |
| No Fear vs CS-Specific Fear | 99.63 |
| No Fear vs Gradient Fear | 1911.1 |
| No Fear vs Overgeneralization | 18,517.7 |

**Table 3.** Model fits for Fear Conditioned Mice. Top table shows the quality of the fit of each model to the behavioral data, expressed as residual sum of squares (RSS, lower shows a better match between model and data), and Akaike Information Criterion (AIC, lower indicates a better match between model and data). Lower table shows the likelihood ratio between the best fitting model (Gradient Fear) and the other models based on their AIC scores.

| Model (Fear Conditioned Group) | RSS | AIC |
| --- | --- | --- |
| Gradient fear | 50.7 | 21.17 |
| Overgeneralization | 69.0 | 23.64 |
| CS-specific fear | 91.7 | 25.91 |
| No Fear | 111.9 | 27.50 |

**Table 3. Model fits for Fear Conditioned Mice.**
| Comparison (Fear Conditioned Group) | Likelihood Ratio (fold likelihood) |
| --- | --- |
| Gradient fear vs Overgeneralization | 3.44 |
| Gradient Fear vs CS-Specific Fear | 10.70 |
| Gradient Fear vs No Fear | 23.71 |

The approach elaborated here focuses on hypothesis testing, in which specific, contrasting models are fully definable *a priori*, such as avoidance of only one odor, or equal avoidance of all odors. However, some models may be hypothesized that depend on incompletely known facts, such as our Gradient Fear model, which estimated a 67% overlap between MV and BA and a 33% overlap between MV and HEX based on neurophysiological data. For some questions, it may be helpful to supplement the hypothesis testing with exploratory data analysis to understand *post hoc* what the optimum model to fit the data would be. This can be performed by iteratively updating the model function to explore the parameter space (and can be automated if using a curve fitting package that supports scripting). To illustrate the point, we note that in our data we found the 67%/33% (MV-BA overlap/MV-HEX overlap) gradient fear model fit 5.2-fold better than a 50%/50% model and 1.46-fold better than an 80%/20% model but was within rounding error of a 75%/25% model.

### Comparison to frequentist analysis

Analyzing data from the arena using traditional frequentist approaches like ANOVA is possible in principle but leads to challenges in concept and interpretation. Conceptually, data from the arena loosely fits with the idea of a two-way, mixed model ANOVA or generalized linear model, where the between-subjects factor is the experimental manipulation (in this case fear conditioning or control exposure) and the within-subjects factor is the arena odor zone (the four corners and four sides), leading to a 2×8 mixed model ANOVA. This is a decidedly imperfect match to the situation, in that the time spent in each zone is not independent of time spent in other zones and our data fails the presumptions of normality and sphericity for the odor zone factor (i.e. the variance of the data is not the same in each odor zone; Mauchly’s Test; W=0.056; p<0.001), but there are no simple alternatives.

Despite these limitations, we used ANOVA to produce a more traditional analysis for comparison to the model testing approach, including performing Greenhouse-Geisser correction of the degrees of freedom to compensate for the non-sphericity of the data. As hypothesized, this analysis shows a very large and statistically significant interaction of fear conditioning by arena zone (F_4.12,42_=18.67, p<0.001, partial eta=0.949). However, this doesn’t demonstrate anything about the relative avoidance of the various odors. That requires *post hoc* testing, which in this design would require 28 pairwise comparisons of odor zones within the fear conditioned group and 28 pairwise comparisons within the control group.

Using the Scheffé *post hoc* test on the controls (which is appropriate for this large number of comparisons), we found no significant differences among any of the odor zones in the control animals.

Using that test on the conditioned animals we found 12 significant differences: time spent in the No Odor corner was greater than in the three other corners and the MV-BA, MV-HEX, and No Odor-BA sides but not the No Odor-HEX side; time spent in the No Odor-Hex side was greater than the MV, BA, and HEX corners and the other three sides. Notably, the choice of *post hoc* test mattered here, as the use of the less selective Bonferroni test (which is more appropriate for cases with small numbers of comparisons) finds a possibly spurious significant difference between the No Odor corner and the side between MV and BA (p=0.031) in the control animals. Taken together, this exercise highlights how the arcana of frequentist statistical analysis can make traditional analyses difficult to apply and interpret even in this simple but multivariate dataset (Cramer *et al*. 2016).

## Discussion

The methods described here allow simultaneous assessment of the approach/avoidance behavior of four odor or control stimuli at the same time. Traditional metrics like distance from a target stimulus or measures of thigmotaxis are readily extracted along with more multivariate information about the overall pattern of the mouse’s avoidance behavior. Finally, we demonstrated a statistically sound method for defining quantitative hypotheses about the shape of that distribution and statistically sound methods for quantifying the relative likelihood of those hypotheses. This method is both efficient and cost-effective, in that it does not require the mouse to be trained in the arena beforehand, lasts only five minutes, and does not require expensive apparatus.

Taken together, these methods produce a very effective means of quantifying how much a mouse generalizes its fear or avoidance response across odors following the traumatic experience of odor-cued conditioning. This is an important case where relatively small differences in the avoidance between odors are needed to distinguish cases like CS-specific fear and fear gradients from potentially disordered cases like overgeneralization across odors. This method has enabled us to recently conclude that GABA_B_ receptor-mediated signaling in the olfactory bulb shapes odor-cued fear generalization such that blockade of GABA_B_ receptors induces overgeneralization while stimulation of GABA_B_ receptors reduces fear generalization (Bakir et al. 2025).

Olfactory science has long struggled to understand smell’s perceptual space, in part because it is difficult to map odors along its disparate perceptual and physical dimensions (Doty 2025; Koulakov *et al*. 2011). While significant progress has been made in human subjects (Lee *et al*. 2023), such perceptual analysis in animal models often requires significant training time relative to the number of datapoints collected (Nakayama *et al*. 2022; Youngentob *et al*. 1990). The present approach has potential value as an efficient means of quantifying perceptual similarities among odors, especially if the analysis were adapted to quantify the best-fitting relationships rather than testing *a priori* models. However, we caution that differences in avoidance do not necessarily correspond to differences in perceptual quality because generalization can also occur along non-sensory, categorical axes (Dunsmoor *et al*. 2011; Gerdes *et al*. 2020; Hanson 1957; 1959). In the demonstration data included here, there is no known reason to expect non-sensory generalization and the pattern of results is consistent with the expectation that an ester is perceptually more similar to another ester than to a ketone. In principle, reaching such a conclusion using this or any other generalization-based assay requires additional experimentation to validate the interpretation of the generalization gradient as a metric of perceptual distance.

We also note that combining this method with electrophysiological or optogenetic manipulations that require surgical implantation introduces tissue damage that could in principle impact olfactory perception directly. In the demonstration data presented here, neither control mice nor MV-cued fear conditioned mice avoided HEX. This presumably indicates a lack of fear generalization from MV to HEX but could in theory result from surgical damage from implanted cannulae that selectively impaired ketone-responsive regions of the bulb. In this case we are confident that this cannula implantation did not impair their perception of HEX because we have previously reported intact discrimination of HEX, MV, and BA in mice implanted with identical cannulae in a spontaneous disinhibition paradigm (Bakir *et al*. 2026). In other studies it may be necessary to perform a similar control experiment or to use a design that permits counterbalancing of which stimulus is reinforced.

We expect most practitioners of this method will find the relative likelihood of competing models and the corresponding graphs to be compelling scientific evidence. However, in some fields it may be that likelihood-based statistics are not sufficient. We thus note the following solutions to specific challenges. First, in contexts where comparison of the observed data to a null hypothesis is required, the inclusion of a No Fear or “chance” model effectively serves as a null hypothesis test. Second, in contexts where one particular zone is central to the interpretation, the hypothesis testing approach illustrated above can complement a more traditional, single-variable test of time spent in that zone via a t-test or Kolmogorov-Smirnov test. Finally, in contexts in which it is desirable to take a frequentist approach to quantify the risk of sampling error, it is possible to define the null hypothesis as equal time in each zone and then use a one-group frequentist statistical test to compare the empirical cumulative distribution function of the data to the theoretical cumulative distribution function of the null or other continuous hypothesis. We recommend the use of the Kuiper test, a non-parametric test similar to the Kolmogorov-Smirnov test, to generate a p-value in this case because a) the data essentially cannot be normally distributed so a non-parametric test is required, and b) the perimeter zones form a circle, which requires a test based on circular distributions (e.g. Kuiper, von Mises, etc.). Finally, to reduce confusion we emphasize that the goal of the function fitting in our approach is not curve-fitting *per se*, where the goal is to change the model to most closely match the data, but only the scaling of the (otherwise unchanged) models to enable optimum quantification of the differences between model and data.

Finally, we note that the positional metrics employed in this demonstration provide a very limited quantification of the complex set of behaviors mice actually engage in during testing. Fear generalization likely also influences behaviors like head orientation, sniffing (Wesson *et al*. 2008), rearing, and whisking. The advent of open-source pose estimation software (e.g. DeepLabCut, FreiPose, and SLEAP) enables the quantification of a much richer set of information about mouse body position from video recordings (Lauer *et al*. 2022; Mathis *et al*. 2018; Pereira *et al*. 2022; Schneider *et al*. 2022; Ye *et al*. 2024) that could reasonably be part of a larger suite of hypotheses tested in the arena. The segmentation of these datasets into behavioral modules (e.g. syllables) with additional analysis tools (e.g. keypoint-MoSeq) would similarly permit a richer interpretation of data from the arena that would complement the basic attraction/avoidance model of animal position described here (Weinreb *et al*. 2024). A challenge for many of these sophisticated behavioral analyses is often the difficulty of interpreting their support for any given experimental hypothesis (Mathis and Mathis 2025). The general approach articulated here, of specifying quantitative hypotheses and computing their relative likelihoods could readily be extended to apply to these more sophisticated behavioral data.

## Acknowledgements

This work was supported by grant R01 MH101293 from the National Institutes of Health. We thank Walt Shotwell and Ben Samuels for their assistance with this project.

## Data Availability Statement

Data available upon request.

